# ProtmRNA: Cross-Modal Knowledge Transfer from Proteins to Messenger RNA

**DOI:** 10.64898/2026.05.19.726141

**Authors:** Gang Xu, Xinyu Wu, Jianpeng Ma

## Abstract

**Motivation:** According to the central dogma of molecular biology, messenger RNA (mRNA) sequences are directly translated into amino acid sequences, positioning mRNA as the fundamental intermediary between genetic information and functional proteins. This natural correspondence suggests that mRNA sequence analysis could greatly benefit from the rich evolutionary and functional representations learned by large-scale protein language models.

**Results:** ProtmRNA repurposes the pre-trained ESM-2 protein language model for mRNA sequence processing via cross-modal transfer learning. Evaluated on mRNA- and protein-related datasets, along with eight additional benchmarks compiled in this study, ProtmRNA achieves performance comparable or superior to state-of-the-art mRNA language models while using less than half the pre-training computational resources. This work establishes the potential of cross-modal transfer learning between biological sequences by demonstrating that protein-derived knowledge can be efficiently transferred to mRNA, offering a resource-efficient paradigm for advancing mRNA sequence understanding.

**Availability and Implementation:** The pre-trained ProtmRNA model and the eight CDS-region regression benchmarks curated in this study are publicly available at https://github.com/pesenteur/ProtmRNA.

## Introduction

In recent years, deep learning has emerged as a transformative force in biological sequence analysis, revolutionizing our ability to decode the complex language of life (Lin, et al., 2023). A wide range of protein language models (PLMs) have been developed to extract rich evolutionary and functional information from amino acid sequences (Xiao, et al., 2025), including the groundbreaking ESM series (Lin, et al., 2023), which has achieved state-of-the-art performance across diverse tasks such as protein structure prediction (Jumper, et al., 2021), variant effect analysis (Zhang, et al., 2024), and protein design (Song, et al., 2025). From a practical standpoint, the availability of pre-trained PLM checkpoints facilitates the rapid development and deployment of specialized models for downstream applications (Hie, et al., 2024; Sun, et al., 2025), offering an efficient and effective solution for biological research. These advances underscore the great promise of extending language model paradigms to other biological sequences, such as RNA (Zhang, et al., 2025) and DNA (Fishman, et al., 2025), ushering in a new epoch of data-driven biological discovery.

Messenger RNA (mRNA) plays a critical role in protein synthesis and gene expression regulation, serving as the essential intermediary that bridges genetic information encoded in DNA to functional proteins. In addition, the unprecedented success of mRNA vaccines in combating infectious diseases has highlighted the urgent need for accurate and efficient computational tools for mRNA sequence analysis (Zhang, et al., 2023). A mature mRNA transcript consists of a coding sequence (CDS) that encodes a protein, flanked by 5’ and 3’ untranslated regions (UTRs), and stabilized by a 7-methylguanosine (m7G) 5’ cap and a 3’ poly(A) tail. In this study, we focus specifically on the CDS region for model training and evaluation, as it is the only region that directly corresponds to amino acid sequences, making it the most natural and biologically meaningful target for cross-modal knowledge transfer from proteins to mRNA.

While early RNA language models were primarily designed for general RNA sequences (Penic, et al., 2025), recent years have witnessed the development of several mRNA-specific models tailored to its unique structural and functional characteristics. CodonBERT (Li, et al., 2024), pre-trained on 10 million full-length mRNA coding sequences from mammals, bacteria, and human viruses using codons as the fundamental input unit, excels at various CDS-related prediction tasks. RNA-FM (Shen, et al., 2024), pre-trained on 45 million mRNA coding sequences, is purpose-built to capture mRNA-specific features and exhibits robust performance on diverse downstream applications. Most notably, mRNABERT (Xiong, et al., 2025), pre-trained on 36 million full-length mRNA sequences, integrates protein sequence information via contrastive learning between CDS embeddings and corresponding amino acid embeddings, achieving state-of-the-art performance in the majority of downstream tasks including 5’ UTR and CDS design, RNA-binding protein (RBP) site prediction, and full-length mRNA property prediction.

In this work, we introduce ProtmRNA, a cross-modal transfer learning framework that repurposes the pre-trained ESM-2 protein language model for mRNA sequence processing. Leveraging the inherent biological correspondence between codons and amino acids defined by the universal genetic code, we extend ESM-2’s 20-amino-acid embedding vocabulary to include 64 codon embeddings. Our experimental results demonstrate that ProtmRNA achieves performance that is either comparable to or surpasses that of state-of-the-art mRNA language models, while consuming less than half the computational resources required to pre-train these models. This work underscores the substantial potential of cross-modal transfer learning in the domain of biological sequence analysis, establishing that knowledge distilled from large-scale protein sequence datasets can be effectively transferred to mRNA sequence understanding.

## Materials and methods

The overall architecture of ProtmRNA is illustrated in Figure 1a. Conceptually, we adopt the pre-trained ESM-2 650M protein language model as the source domain for cross-modal knowledge transfer. We extend ESM-2’s original 20-amino-acid embedding space to incorporate 64 codon embeddings, where each codon embedding is initialized to match the embedding of its corresponding amino acid according to the universal genetic code. The original ESM-2 vocabulary contains 33 tokens (20 standard amino acids plus 13 special tokens). We replace the 20 amino acid tokens with 64 codon tokens and add one additional token (“XXX”) to represent unknown codons, resulting in a final vocabulary size of 78 tokens. The hidden dimension of ProtmRNA remains identical to that of ESM-2 at 1280, preserving both the model’s original architectural capacity and the rich evolutionary knowledge encoded during its pre-training.

**Figure 1.**
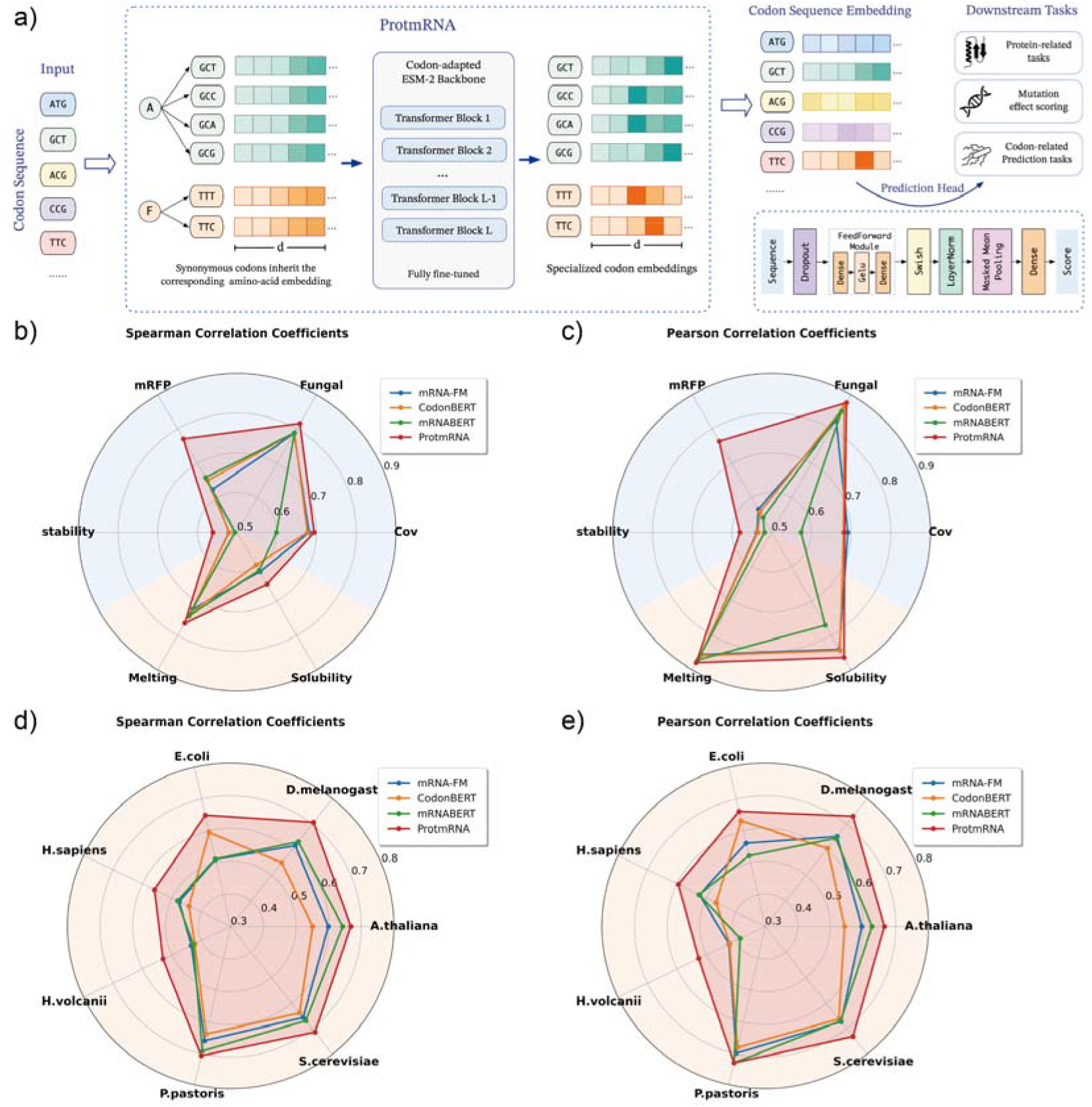
a) Overview of the ProtmRNA framework. b, c) Spearman and Pearson correlation coefficients of different mRNA language models on diverse downstream tasks. d, e) Spearman and Pearson correlation coefficients of different mRNA language models on transcript abundance prediction across seven species. CDS-related and protein-related tasks are highlighted with blue and orange backgrounds, respectively.

For pre-training, we use a subset of the mRNABERT training dataset (Xiong, et al., 2025), randomly sampling 4.8 million full-length mRNA coding sequences with a maximum token length of 768. Pre-training is performed on 4 Nvidia V100 GPUs with a batch size of 24 per GPU, for a total of 50,000 training steps. We use the AdamW optimizer with an initial learning rate of 1e-4. All pre-training experiments are implemented in TensorFlow 2.4.

To reduce computational overhead and ensure fair comparison across methods, we freeze the pre-trained mRNA language model during downstream evaluation and use it to extract fixed-length representations for all mRNA sequences in the benchmark datasets. This approach is practically justified because in many real-world scenarios, only pre-extracted representations from language models are used for downstream tasks. Although joint fine-tuning of the large language model and downstream task head may yield better performance, it is often impractical for multi-modal tasks or tasks with very large datasets. Identical downstream task heads are used for all compared methods (Figure 1a). For downstream training, we use the Adam optimizer with an initial learning rate of 1e-3, which is reduced by half when the validation accuracy decreases. Training is run for a maximum of 100 epochs, with early stopping triggered if the learning rate is reduced 5 times in total. The batch size is set at 8 for all tasks.

All benchmarks used in this study are regression tasks. We therefore use two standard correlation metrics for performance evaluation: Spearman correlation coefficient and Pearson correlation coefficient.

## Results

For CDS-related downstream tasks, we evaluate ProtmRNA and other codon-based models on four regression tasks: SARS-CoV-2 vaccine degradation prediction (Wayment-Steele, et al., 2022), fungal gene expression prediction (Nasr, et al., 1996), mRFP expression prediction (Nieuwkoop, et al., 2023), and mRNA stability prediction (Medina-Muñoz, et al., 2021). For protein-related tasks, we include two regression tasks: protein melting point prediction and protein solubility prediction (Xiong, et al., 2025). We also adopt the transcript abundance prediction benchmark across seven species from Xiong et al. (Xiong, et al., 2025). As shown in Figure 1b-e, ProtmRNA achieves performance comparable or superior to other leading mRNA language models. We further compare ProtmRNA with its starting checkpoint, ESM-2 650M, in Table S1-2. The results show that ProtmRNA outperforms ESM-2 in most cases, even on protein-related tasks, confirming that our cross-modal knowledge transfer strategy enhances mRNA sequence understanding while preserving protein-related predictive capability.

In addition, we compile eight CDS-region regression benchmarks from high-quality datasets in the MaveDB repository (Table S3). All source records are converted into a standardized sequence-label format, where each entry consists of a full-length CDS sequence as input and its corresponding continuous experimental score as the label. Each benchmark is split into training, validation, and test sets using a fixed random seed. These eight benchmarks span diverse biological functions, including enzymatic activity, membrane protein expression, and proteolysis-based folding stability, ensuring a comprehensive evaluation of model generalization across different biological contexts. As shown in Table S4, ProtmRNA also achieves better results than other leading mRNA language models in most cases. These results demonstrate that ProtmRNA generalizes better to protein-related tasks than existing mRNA language models, likely due to the rich functional and evolutionary representations inherited from the ESM-2 protein language model.

## Concluding discussion

The central finding of this study is that evolutionary and functional knowledge encoded in large-scale protein language models can be efficiently transferred to mRNA sequence analysis through a biologically inspired cross-modal transfer learning framework. Our results demonstrate that ProtmRNA, built by repurposing the pre-trained ESM-2 model with codon-adapted embeddings, achieves competitive or superior performance compared to state-of-the-art mRNA-specific language models across a comprehensive set of benchmark tasks, while requiring less than half the computational resources for pre-training. This success validates our core hypothesis that the intrinsic biological correspondence between codons and amino acids, as defined by the universal genetic code, provides a natural and robust bridge for knowledge transfer between these two fundamental biological modalities.

A key advantage of ProtmRNA over existing mRNA language models lies in its ability to leverage decades of accumulated progress in protein language modeling. Unlike previous approaches that either pre-train mRNA models from scratch (CodonBERT, RNA-FM) or integrate protein information through auxiliary contrastive learning objectives (mRNABERT), our framework directly inherits the rich evolutionary representations learned by ESM-2 from hundreds of millions of protein sequences. This not only drastically reduces the computational burden of model development, but also ensures that ProtmRNA captures deep functional and structural information that is inherently linked to mRNA sequences through the translation process. Notably, ProtmRNA’s strong performance on both mRNA-related tasks and protein-related tasks further confirms that mRNA sequences contain implicit information about the structure and function of their encoded proteins, and that our cross-modal approach effectively extracts this cross-talk information.

Another important contribution of this work is the compilation of eight new CDS-region regression benchmarks from the MaveDB repository. Existing mRNA sequence analysis benchmarks are often limited to a narrow range of tasks, such as stability prediction or gene expression estimation. In contrast, our curated benchmarks span diverse biological functions including enzymatic activity, membrane protein expression, and proteolysis-based folding stability, providing a more comprehensive evaluation framework for future mRNA language models. These benchmarks will facilitate the development of models that can accurately predict the functional consequences of mRNA sequence variations, which is critical for applications such as mRNA vaccine design and therapeutic protein engineering.

Looking beyond mRNA, the cross-modal transfer learning paradigm introduced in this work has broad implications for biological sequence analysis in general. Many biological molecules share intrinsic functional and evolutionary relationships, and knowledge transfer between these modalities could significantly accelerate the development of models for data-scarce sequence types. For instance, similar strategies could be applied to transfer knowledge from proteins to tRNA or rRNA, or from mRNA to genomic DNA, representing promising directions for future work.

Beyond the benchmarks presented here, mRNA language models hold significant practical value across multiple domains of RNA biotechnology. In protein engineering, they can guide synonymous codon optimization to enhance folding stability without altering the protein sequence. For recombinant expression, accurate models enable the design of high-yield mRNA constructs by predicting transcript stability and translational output directly from sequence. Most notably, in mRNA vaccine development, sequence-level models offer a rapid computational route to optimizing vaccine candidates for degradation resistance, translational efficiency, and reduced immunogenicity. As mRNA therapeutics continue to expand beyond infectious diseases into cancer and rare genetic disorders, robust and resource-efficient mRNA language models will become essential tools for accelerating candidate design and ensuring therapeutic efficacy.

## Acknowledgements

J.M. wants to thank the support from the National Key Research and Development Program of China (No. 2024YFA1307502), the Science and Technology Innovation Plan of Shanghai Science and Technology Commission (No. 23JS1400200), the Research Fund for International Senior Scientists (No. W2431060), and the Research Fund of Development and Reform Commission of Hunan province (No. 2507-430000-04-05-666048). G.X. wants to thank the support from the National Natural Science Foundation of China (No. 32300535).

## Author Contributions

G.X. developed and trained the model. X.W. collected the data and performed feature extraction. G.X. and X.W. conducted the downstream task evaluations. G.X. and X.W. drafted the manuscript. J.M. and G.X. supervised the research and designed the study. All authors contributed to project discussions and manuscript revisions.

## Notes

### Competing Interest Statement

The authors have declared no competing interest.

