## Supplementary material for "ProtmRNA: Cross-Modal Knowledge Transfer from Proteins to Messenger RNA": SI

Xu *et al*.

**Table S1.** Spearman correlation coefficients of ProtmRNA and ESM-2 on CDS-related and protein-related downstream tasks.

|  | Cov | Fungal | mRFP | stability | Melting | Solubility |  |
| --- | --- | --- | --- | --- | --- | --- | --- |
| ESM-2 | 0.552 | 0.731 | - | 0.514 | 0.649 | 0.666 |  |
| ProtmRNA | 0.694 | 0.816 | 0.771 | 0.561 | 0.764 | 0.651 |  |
|  | A.thaliana | D.melanogaster | E.coli | H.sapiens | H.volcanii | P.pastoris | S.cerevisiae |
| ESM-2 | 0.592 | 0.654 | 0.609 | 0.504 | 0.457 | 0.670 | 0.679 |
| ProtmRNA | 0.668 | 0.707 | 0.649 | 0.559 | 0.530 | 0.706 | 0.714 |

**Table S2.** Pearson correlation coefficients of ProtmRNA and ESM-2 on CDS-related and protein-related downstream tasks.

|  | Cov | Fungal | mRFP | stability | Melting | Solubility |  |
| --- | --- | --- | --- | --- | --- | --- | --- |
| ESM-2 | 0.540 | 0.765 | - | 0.539 | 0.818 | 0.767 |  |
| ProtmRNA | 0.684 | 0.877 | 0.764 | 0.580 | 0.880 | 0.865 |  |
|  | A.thaliana | D.melanogaster | E.coli | H.sapiens | H.volcanii | P.pastoris | S.cerevisiae |
| ESM-2 | 0.591 | 0.680 | 0.628 | 0.565 | 0.472 | 0.674 | 0.700 |
| ProtmRNA | 0.665 | 0.731 | 0.661 | 0.595 | 0.526 | 0.729 | 0.731 |

**Table S3.** Details of the eight CDS-region regression benchmarks compiled in this study.

| **Name** | **Description** | **Train** | **Val** | **Test** |
| --- | --- | --- | --- | --- |
| PTEN | PTEN variant scores from a lipid phosphatase activity assay in a humanized yeast model. This dataset provides a disease-relevant enzymatic activity regression target for CDS-level sequence-to-score prediction. | 1575 | 394 | 394 |
| psaE | Photosystem I reaction center subunit IV (psaE) variants measured by cDNA-display proteolysis using trypsin and chymotrypsin. The regression label is the coupled-model ΔG value, reflecting protein folding stability. | 2966 | 742 | 742 |
| Plcg1 | Phospholipase C gamma 1 (Plcg1) SH3-domain variants measured by cDNA-display proteolysis using trypsin and chymotrypsin. The regression label is the coupled-model ΔG value, reflecting folding stability of a signaling-related protein domain. | 3000 | 750 | 750 |
| MJ1198 | Uncharacterized protein MJ1198 variants measured by cDNA-display proteolysis using trypsin and chymotrypsin. The regression label is the coupled-model ΔG value, providing a stability-focused dataset from an archaeal protein target. | 2924 | 731 | 731 |
| HECTD1 | HECTD1 variants from a trypsin-digestion cDNA-display proteolysis experiment. The dataset is used as a folding-stability regression benchmark for a ubiquitination-related protein target. | 5376 | 1345 | 1345 |
| UBE4B | UBE4B variant score set included as a continuous CDS-level sequence-to-score regression dataset. Biologically, UBE4B is a ubiquitination/proteostasis-related protein; the description is kept conservative because the score-set metadata should be cited directly from MaveDB before claiming a specific protease protocol. | 3209 | 803 | 803 |
| VLN4 | Villin-4 (VLN4) variants measured by cDNA-display proteolysis using trypsin and chymotrypsin. The regression label is the coupled-model ΔG value, representing folding stability for a plant actin-bundling protein domain. | 3044 | 762 | 762 |
| JAG1 | JAG1 variant scores from a membrane-expression assay using a cDNA variant library. This dataset provides a disease-relevant functional-expression regression target associated with Alagille syndrome variant interpretation. | 1888 | 472 | 472 |

**Table S4.** Results of codon-based RNA language models on the eight compiled datasets.

| Spearman | PTEN | psaE | Plcg1 | MJ1198 | HECTD1 | UBE4B | VLN4 | JAG1 |
| --- | --- | --- | --- | --- | --- | --- | --- | --- |
| mRNA-FM | 0.298 | 0.752 | 0.579 | 0.752 | 0.818 | 0.686 | 0.628 | 0.226 |
| CodonBERT | 0.048 | 0.835 | 0.682 | 0.842 | 0.901 | 0.805 | 0.764 | 0.321 |
| mRNABERT | 0.151 | 0.741 | 0.651 | 0.708 | 0.767 | 0.566 | 0.504 | 0.183 |
| ProtmRNA | 0.393 | 0.913 | 0.831 | 0.940 | 0.941 | 0.895 | 0.792 | 0.356 |
| Pearson | PTEN | psaE | Plcg1 | MJ1198 | HECTD1 | UBE4B | VLN4 | JAG1 |
| mRNA-FM | 0.310 | 0.689 | 0.573 | 0.713 | 0.804 | 0.623 | 0.542 | 0.311 |
| CodonBERT | 0.180 | 0.775 | 0.660 | 0.800 | 0.897 | 0.735 | 0.827 | 0.352 |
| mRNABERT | 0.109 | 0.673 | 0.630 | 0.667 | 0.747 | 0.500 | 0.494 | 0.254 |
| ProtmRNA | 0.355 | 0.856 | 0.847 | 0.888 | 0.943 | 0.812 | 0.783 | 0.342 |
